# Global Convergence of Plant Functional Trait Composition in the Anthropocene

**DOI:** 10.64898/2026.03.27.714725

**Authors:** Sophie Wolf, Daria Svidzinska, David Schellenberger Costa, Miguel D. Mahecha, Julia Joswig, Lirong Cai, Karin Mora, Guido Kraemer, Kolja Nenoff, Marten Winter, Christian Wirth, Susanne Tautenhahn, Helge Bruelheide, Mark van Kleunen, Holger Kreft, Petr Pyšek, Patrick Weigelt, Teja Kattenborn

## Abstract

Since the onset of European colonial expansion, humans have accelerated species migration across continents, reshaping plant functional composition and associated ecosystem processes. Plant functional traits–such as leaf area, plant height, or rooting depth–are structured along major axes of variation, including size and leaf economics, that reflect ecological strategies. While human-mediated changes in this trait space have been documented regionally or for specific taxa, there exists no global, grid cell-level quantification of past shifts across major axes of trait variation. Here, we link global citizen science plant occurrence data with data on 37 above- and below-ground traits, and information on native and introduced status for each occurrence. Using dimension-reduction on grid cell-level trait means and introduced species status as a proxy for anthropogenic change, we identify three major axes of functional variation: the size, leaf economics, and life-span axes. By comparing past (native-only) and present-day trait distributions in 3D trait space and geographically, we find prominent region-specific shifts along all three axes. Overall, functional composition converges toward (mostly) smaller, more acquisitive, and shorter-lived assemblages, with region-specific differences in which axis shifts are most pronounced. These results provide the first global estimate of how human-mediated plant introductions have altered ecosystem functional composition in the past centuries, highlighting the spatial patterns and trait dimensions most affected by anthropogenic pressures.

## Introduction

Throughout human history, and especially since the onset of European colonial expansion, humans have profoundly reshaped global plant biogeography (Capinha et al, 2015). Although the Anthropocene is not uniformly defined, one proposal places its onset at this collision of the so-called Old and New Worlds between 1492 and 1800 (Lewis and Maslin, 2015), a period marked by unprecedented human-mediated species migration across continents, large-scale range modifications, and the homogenization of once distinct biota (Winter et al, 2009; Yang et al, 2021; Holmes et al, 2024). These biogeographic changes have often resulted in shifts in community composition, which are often driven by anthropogenic environmental changes such as land-use change (logging, urbanization, etc.) and–more recently–climate change (Lenzner et al, 2022; Liu et al, 2023, 2024; Nagy et al, 2021; Tautenhahn et al, 2026).

Terrestrial ecosystems are largely structured by plants, which dominate biomass and shape habitat structure (Bar-On et al, 2018; Scheiter et al, 2024). Plant functional traits—such as plant height or leaf nitrogen content—mediate energy and nutrient fluxes and reflect life-history strategies (Violle et al, 2007; Divǐsek et al, 2018, 2025; Guo et al, 2022, 2018; Lavorel and Garnier, 2002). Consequently, shifts in plant community composition and thus in functional composition can be expected to have cascading ecosystem consequences (Wirth and Lichstein, 2009; Chapin et al, 2000; Reichstein et al, 2014). Many of these plant functional traits are highly correlated, and previous work has shown that both species and community strategies are structured along general axes of variation within high-dimensional trait space (Díaz et al, 2016; Bruelheide et al, 2019; Weigelt et al, 2021; Joswig et al, 2022; Reich, 2014). This “global spectrum of plant form and function” defines interpretable axes—size variation and leaf economics variation—along which ecosystem functional composition can be studied and compared in relation to environmental properties (Joswig et al, 2022; Sun et al, 2026).

Several recent studies have documented shifts in plant functional composition at local to regional scales using re-survey vegetation plot data, often indicating trait convergence or homogenization and shifts toward more acquisitive strategies (Klinkovská et al, 2025; Aguirre-Gutiérrez et al, 2020; Niu et al, 2016). Other approaches found similar shifts in plant functional traits following extinction and introduction of new species at a global scale for tree species (Guo et al, 2026) and in vegetation plots along a gradient of introduced species (Garbowski et al, 2024). However, these approaches focus on a limited number of species or regions, and there is no global assessment of how human-mediated pressures have altered plant functional composition. Specifically, there exists no biogeographic mapping of functional changes since the onset of the Anthropocene at grid-cell level and quantification of shifts across multiple trait axes.

This research gap partly reflects limited biodiversity data, both on plant distributions, and on their temporal changes. Citizen science datasets now provide millions of opportunistically sampled occurrence plant records worldwide (GBIF, 2025), which can capture biogeographic patterns of species composition (Mahecha et al, 2021) and, when linked to trait data, reconstruct robust functional trait distributions at the grid-cell-level (Wolf et al, 2022; Lusk et al, 2026; Mora et al, 2024). To address the lack of temporally resolved plant occurrence data, we propose using introduced species occurrences as indicators of anthropogenic change. Introduced species often co-occur with other human pressures, such as land-use and climate change, rather than acting as primary drivers themselves (Jaureguiberry et al, 2022; Jauni et al, 2015; Ricotta et al, 2009; Pyšek et al, 2005), yet previous work has identified introduced species as the main drivers of changes in functional trait composition specifically (Garbowski et al, 2024; Jaureguiberry et al, 2022).

Here, we combine global opportunistically sampled citizen science plant occurrence data with species level trait information from TRY (Kattge et al, 2020; Joswig et al, 2023), using introduced species occurrences as a proxy for human-mediated changes in functional composition at two time steps: before and after the onset of European colonial expansion, which we will refer to as *past* and *present* for simplicity. Trait values are aggregated into grid-based means, which are then projected into a dimension-reduced global trait space. This allows us to identify major axes of trait variation, and quantify shifts in functional composition both within trait space and across biogeographic regions. Our approach provides the first global, grid-cell-level (approx. 1.5°) estimates of human-mediated changes in plant functional composition along major trait axes, capturing regional convergence and divergence and setting the stage for a spatially explicit understanding of anthropogenic impacts.

## Results

### Global structure of plant functional trait space

To characterize global plant functional trait variation, we aggregated 63 million openly available citizen science (CS) plant occurrence data, linked to 42,520 species trait averages from gap-filled TRY data (see 14), into grid-based means. Grid-cell means were weighted by the number of observations per species in each grid cell, building on Wolf et al (2022); Lusk et al (2026). Each grid cell was represented twice: once including all species (present) and once including only native species (proxy for past grid-cell mean, based on species’ native/introduced status from WCVP and GloNAF (Gov, 2025; Davis et al, 2025), see 14). We then conducted a principal component analysis (PCA) on all grid-cell means, covering 37 traits and both time steps.

A PCA over 37 traits yielded three main axes of functional variation globally. Principal component 1 (PC1) captures the axis of size traits (small-large) and explains 35% of the variation, with traits such as plant height and seed length contributing to this dimension (Fig. 1a,c,d). PC2 (17%) reflects variation in traits related to plant leaf and stem economics (conservative-acquisitive), which is reflected by high specific leaf area (SLA) and high leaf phosphorus (P) on the one hand and high leaf thickness, leaf nitrogen (N) per area, and stem specific density (SSD) on the other end of the spectrum (Fig. 1b,c,d). Leaf size traits (e.g., leaf area, leaf width) contribute to both PC1 and PC2 (see Fig. 1a,d). PC3 explains 8% of the variance and reflects traits related to variation in reproductive strategy and lifespan (fast–slow life-history), with high leaf N (per mass and per area), leaf delta 15N, and germination efficiency on the one hand and high wood ray density, and wood vessel element length on the other end of this dimension (Fig. 1b,c,d). From PC4 onward, the variance explained levels off (see Supplemental Fig. 6).

**Fig. 1:**
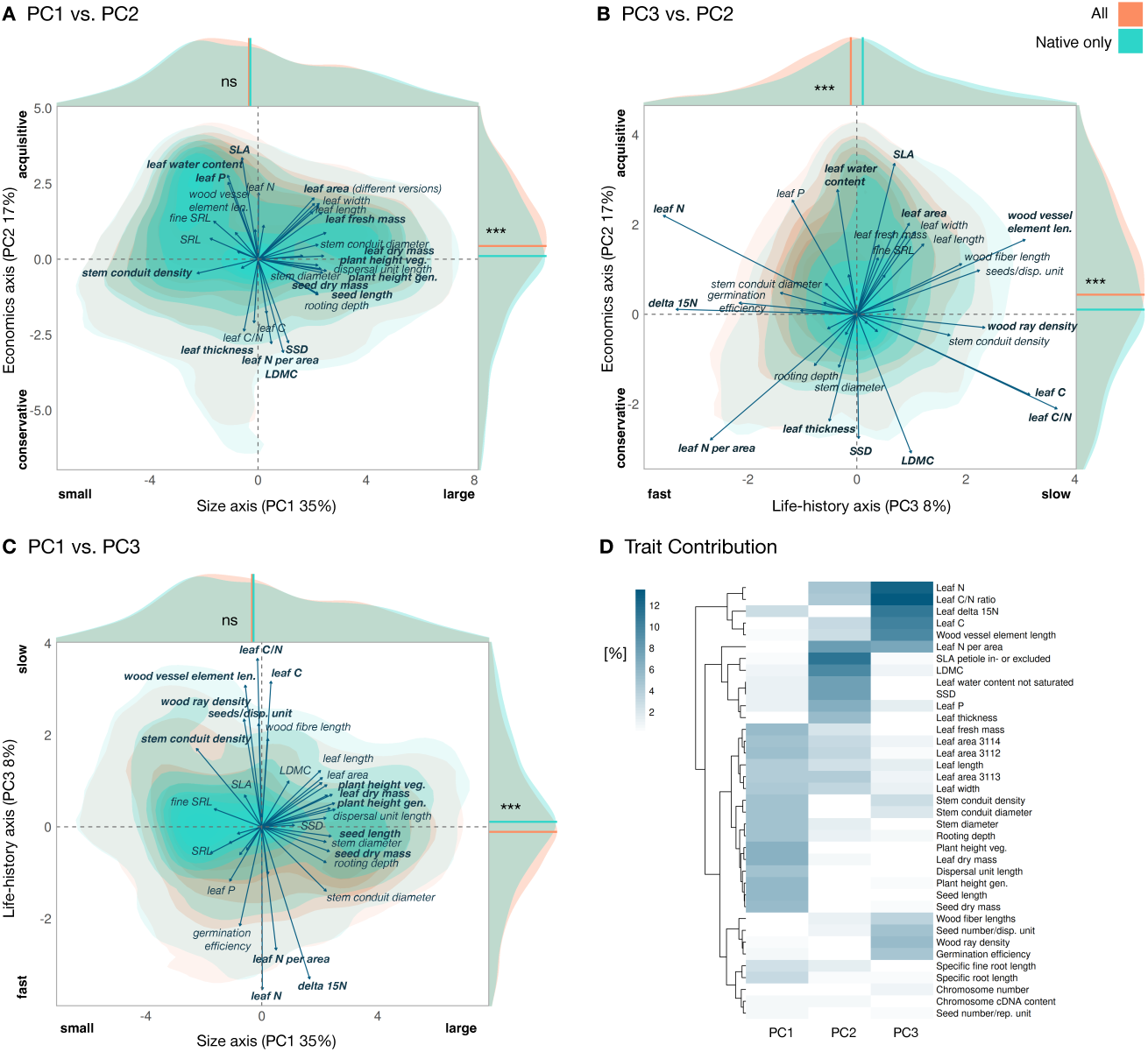
**A-C** Biplots of the first three PCA axes, derived from 37 trait grid-cell means for two time steps (past and present). Traits contributing more than 5% to an axis are labeled in bold in the respective biplot. Orange and mint indicate density estimates for present and past grid-cell averages, respectively. **D** Trait contributions to each of the first three principal components, with hierarchical clustering of traits.

The global present-day biogeography of PC1, PC2, and PC3 (Figure 2) reveals broad-scale patterns, despite noise in regions with sparse data. PC1 follows a temperature gradient across both latitude and elevation (Figure 2a). PC1 scores tend to be larger closer to the equator and decrease toward the poles and at higher elevations, such as in the Andes and the Himalaya. PC2 reflects patterns of climatic seasonality (Figure 2b). Especially in harsh environments (e.g., deserts, alpine regions) PC2 scores are low. In contrast, in seasonally variable environments (e.g., deciduous temperate forests or seasonal rainforests) that rely on acquisitive leaf economics, PC2 scores are high. The third axis (PC3) reflects the high proportion of annual species in desert regions as well as temperate and boreal forested ecosystems at the other end of the spectrum (Fig. 2c).

**Fig. 2:**
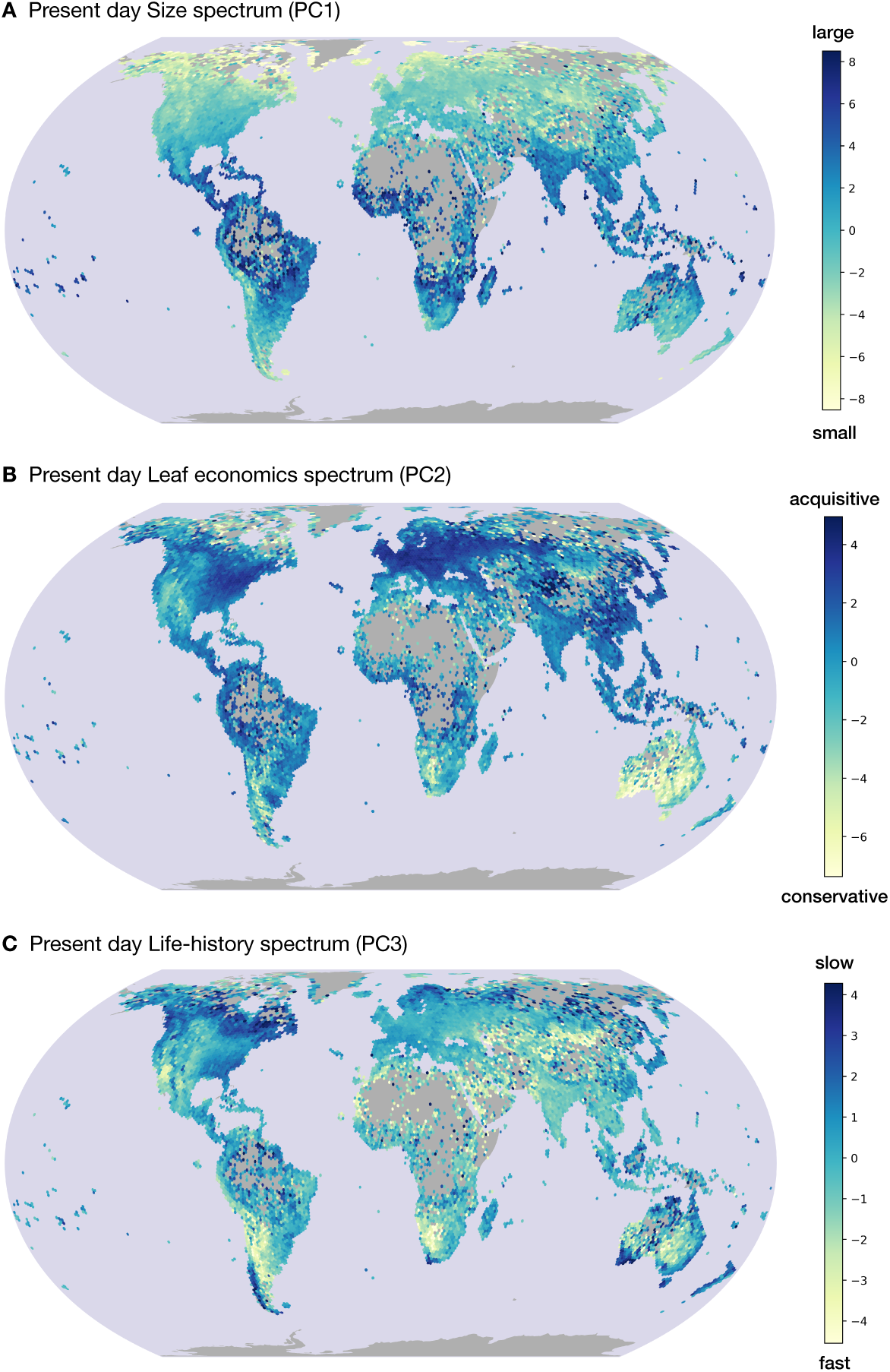
Maps show the present-day spatial distribution of the first three PCA axes derived from 37 above- and below-ground trait grid-cell averages, weighted by the number of observations per species in each grid cell. **A** Size axis, **B** Economics axis, and **C** Life-history axis. Values are unitless PCA scores for each grid cell.

### Global and regional shifts in functional trait composition

Comparing the densities of past and present grid-cell means in trait space shows that including introduced species shifts the distribution of grid-cell averages within the global trait space (Fig. 1). At the global scale, median PC scores shift significantly along PC2 and PC3 (p *<* 0.001), while no significant shift is observed along PC1 (p = 0.19).

The proportion of introduced species occurrences per grid cell in the CS data varies across regions and shows distinct global patterns (Fig. 3). These patterns likely reflect both ecological variation and regional differences in sampling intensity and bias. Particularly high proportions of introduced species occurrences are found across Indomalaya (India, Sri Lanka, and Southeast Asia), exceeding beyond 60% of CS observations in some grid cells. Additional potential hotspots include Buenos Aires Province (Argentina) and parts of West Africa along the Gulf of Guinea. A correlation with human population density (log10) yields Pearson’s *r* = 0.52 and a p-value *<<* 0.01 (see Supplemental Fig. 7). The proportion of introduced species occurrences does not correlate with observation density (log10) per grid cell (*r* = 0.02, see Supplemental Fig. 8). High proportions are also evident on several islands and oceanic archipelagos, including Hawaii, the Azores, and Mauritius.

**Fig. 3:**
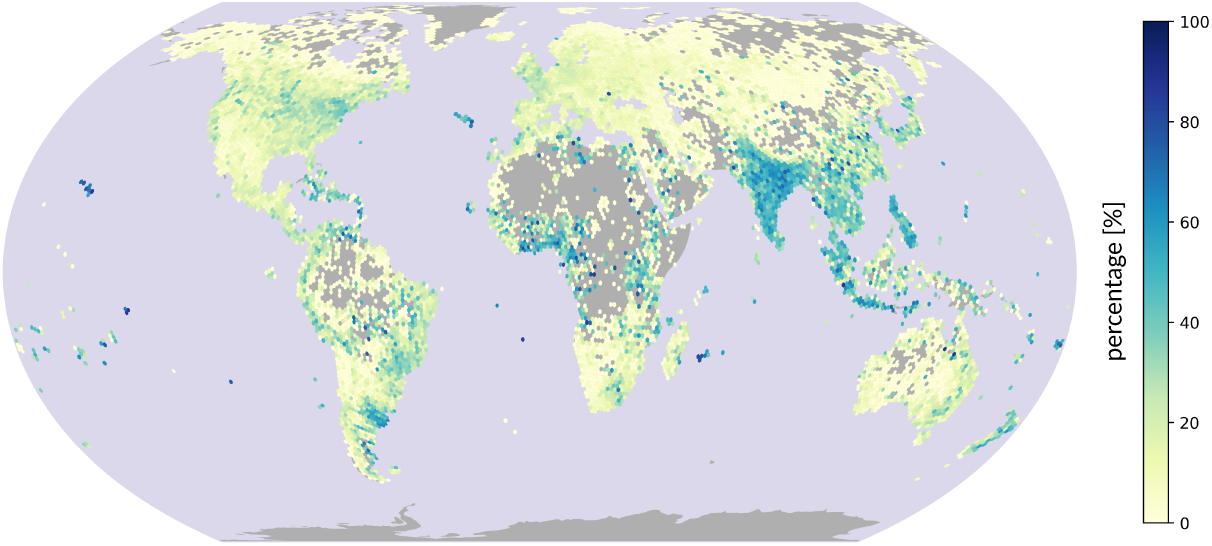
Percentage of citizen science (CS) introduced species observations per grid cell.

To evaluate regionally varying effects of these introduced species occurrences on functional trait composition, we calculated the difference for each grid cell in PCA space along the first three axes (Fig. 4). Although no global shift was detected along the size axis (PC1) (Fig. 1a), grid-level differences reveal regionally opposing patterns: PC1 shifts toward larger grid-cell means in Europe and toward smaller grid-cell means in Eastern North America, India, and Southeast Asia (Fig. 4a). Along the economics axis (PC2), many regions—including Western North America, South America, Southern Africa, India–Southeast Asia, and Australia–New Zealand—shift toward more acquisitive strategies (higher PC2 scores; Fig. 4b). Shifts along PC3 trend toward smaller scores (shorter-lived, faster strategies), with the largest changes in North America, South Asia, New Zealand, and Southern Africa. Many regions shifted along two or even all three axes. We identified regions with prominent shifts using Mann–Whitney U tests with Benjamini–Hochberg–adjusted *p ≤* 0.05 and effect size Cliff’s delta *>* 0.11 (see Fig. 4a–c for regions where median shifts met these thresholds).

**Fig. 4:**
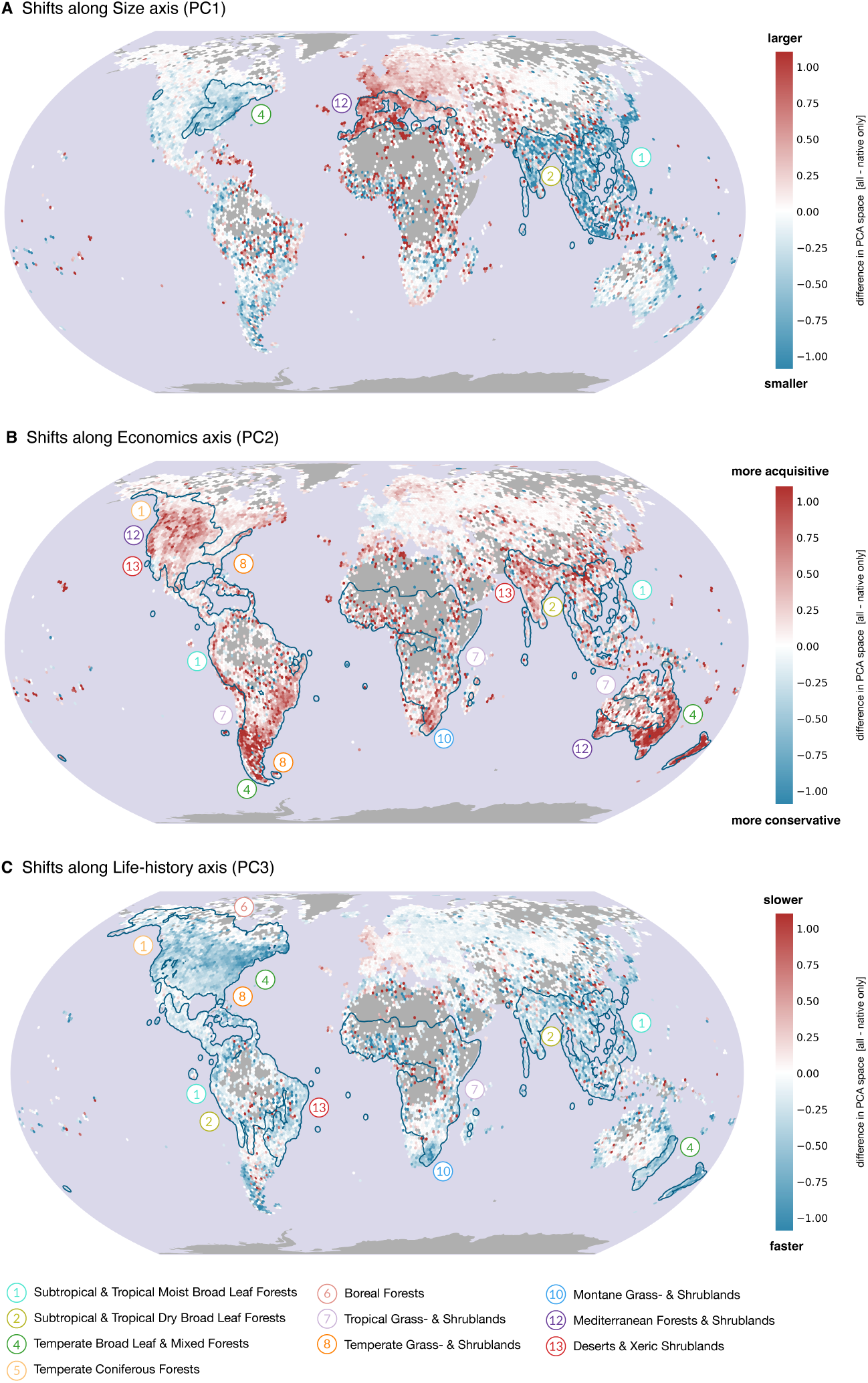
Difference maps showing shifts in trait space along the first three PCA axes: **A** Size, **B** Economics, and **C** Life-history. The outlines indicate biogeographic regions (WWF biomes split by continental realms) with significant overall differences (unpaired Mann-Whitney U test, adjusted *p ≤* 0.05 and deffect size Cliff’s *δ ≥* 0.11). Adjacent polygons were dissolved for visual clarity; number labels indicate the regions; full polygon shapes are shown in Fig. 5.

In addition to quantifying the shifts in PCA space, we calculated significance and effect sizes for each of the 37 traits separately in each region. Effect sizes for each trait and region show that leaf N, leaf P, leaf dry matter content (LDMC), specific root length (SRL), germination efficiency, and SLA are among the traits that shift most often across regions (Supplemental Fig. 9).

### Convergence of regional functional composition

To assess how regions shift within the global PCA space in relation to each other, we plotted the regional centroid shifts (start: centroid of past grid-cell means, end: centroid of present day grid-cell means) for regions with the most prominent shifts along either PC1, PC2, and/or PC3 (see 14). A strikingly consistent pattern emerged, showing a general directionality of regional shifts within trait space (see Fig. 5) converging towards more acquisitive and faster, and often smaller trait composition in this three-dimensional trait space.

**Fig. 5:**
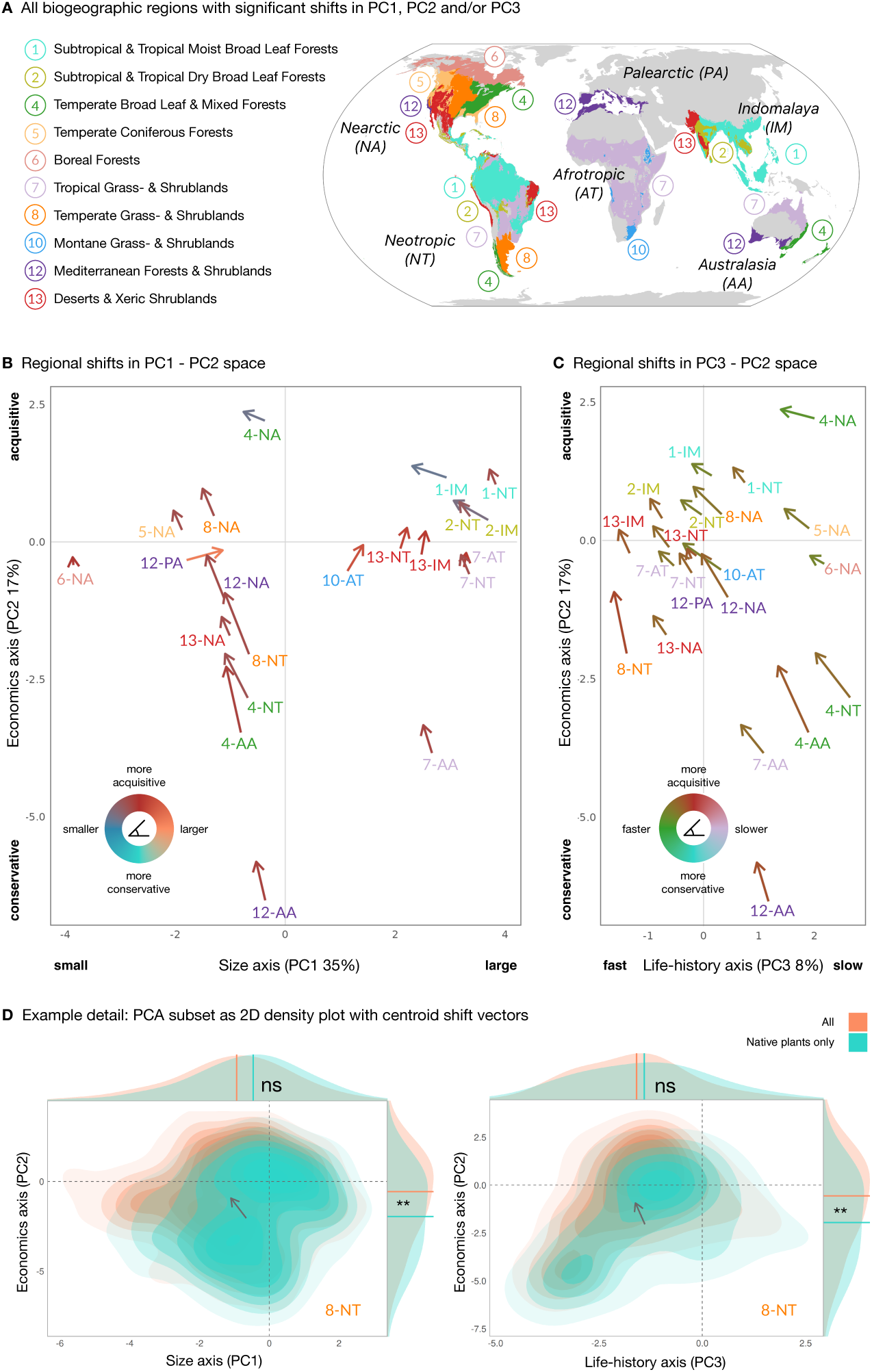
Significant regional shifts in trait space. **A** Regions exhibiting significant median shifts along PC1, PC2, and/or PC3 (Man-Whitney-U test with Benjamini-Hochberg-corrected p-values *<* 0.05 and effect sizes Cliff’s delta *≥* 0.11). **B–C** Vectors from centroids of regional past to present grid-cell mean PCA scores, with vector colors indicating the direction of the shift. **D** Example detail of regional density plots with centroid shift vectors illustrating the methodology.

The direction of shifts depends on the initial position of a region in trait space (see vector plot Fig. 5b). In some regions, shifts are strongest along the leaf economics spectrum (PC2), such as in the Temperate Broadleaf Forests of the Neotropics and Australasia. In contrast, regions already located at the more acquisitive end of the global trait PCA—such as the Tropical Dry and Moist Broadleaf Forests of Indomalaya and the Temperate Broadleaf Forests of the Nearctic—show the strongest shifts along the size axis (PC1), with grid-cell means moving toward smaller values. The Palearctic Mediterranean region is the only one shifting toward larger PC1 values. Similarly, shifts in PC2/PC3 space converge toward more acquisitive and faster positions in trait space (Fig. 5c). Regions at the conservative end mainly shift along PC2, whereas already acquisitive regions shift along PC3 toward shorter-lived grid-cell means.

## Discussion

We characterized the global biogeography of plant functional traits in three primary dimensions. Within this global 3D trait space, we find that regional plant functional composition seem to have undergone convergent shifts since the onset of the Anthropocene. Our results reveal functional composition shifting toward more acquisitive leaf and shorter-lived whole plant strategies when introduced species occurrences are included (see Fig. 5). Shifts in size-related traits show opposing trends when comparing Europe to the rest of the world with Europe shifting towards larger trait means, while many other regions shift towards the smaller end of the size axis. Regional differences in the direction and magnitude of these shifts likely reflect environmental context (initial position in trait space) and human influence.

Building on the “Global Spectrum of Plant Form and Function” (Díaz et al, 2016; Bruelheide et al, 2019; Weigelt et al, 2021), i.e., reducing global plant functional trait variation to its key dimensions, we provide a framework that allows us to quantify regional shifts, compare them across space, and detect convergence or divergence systematically, revealing patterns that would otherwise be obscured in the full high-dimensional trait space (37 traits). Our results reveal three major dimensions of plant functional composition world-wide: Principal components (PCs) 1 and 2 (see Fig. 1a) mirror axes of trait variation that have been previously described both on a species-level (Díaz et al, 2016; Weigelt et al, 2021) and for plant communities (Bruelheide et al, 2019). These two axes have been interpreted as the size axis (small-large), and the leaf and stem economics axis (conservative-acquisitive) (Wright et al, 2004; Zhao et al, 2017), reflecting plant structural investment and leaf robustness.

While PC1 and PC2 were assessed in several studies (e.g., Díaz et al (2016); Wright et al (2004); Bruelheide et al (2019) across species and at the community level, there was so far no consistent and large scale data-driven assessment of a third axis. In our results, PC3 shows a clear biogeographic pattern (Fig. 2) and the contributing traits are high leaf nitrogen (N) (both per mass and per area), germination efficiency and leaf *δ*15N on the one hand, and high carbon (C) content, wood vessel element length, and wood ray density on the other (Fig. 1). We find that PC3 represents a trait dimension that differentiates between nitrogen-rich, short-lived strategies from long-lived strategies, independent of plant height. This interpretation is supported by the correlations with the proportion of annual species occurrences (r = -0.34) and conifer occurrences (in the regions where they exist)(r = 0.39). The proportion of woody species occurrences only has a weak correlation with PC3 (r=0.10) (see Supplemental Fig. 10). This interpretation of the empirically derived PC3 axis mirrors long-established ecological theory of plant strategies, in which a third axis describes ruderal life-history strategies (Grime, 1988; Pierce et al, 2017b; Liu et al, 2025a). Our results therefore align with both data-driven and theory-based frameworks describing major axes of plant functional variation (Díaz et al, 2016; Bruelheide et al, 2019; Grime, 1988; Liu et al, 2025a; Pierce et al, 2017a).

Regional centroid shifts within this global trait space display directionality, i.e., an apparent convergence towards the “small, fast, and short-lived” quadrant in 3D trait space (see Fig. 5). The convergence towards this region in trait space occupied by native European assemblages may partly reflect the legacy of European colonialism. Previous work has shown that European plants contribute disproportionately to the global introduced flora, and that species composition is often more similar among regions that were historically occupied by the same European country (van Kleunen et al, 2015; Lenzner et al, 2022). This opposing shift is particularly pronounced along the size axis (Fig. 4). The observed overall convergence may also reflect adaptation to similar anthropogenic pressures, as plants in early successional stages following disturbance are known to exhibit comparable trait profiles. Together, these patterns suggest potentially additive, and possibly interactive, effects of introduction history and land-use change.

Although our approach is based only on grid-cell means, as opposed to other functional diversity metrics, the converging shifts toward a common region of trait space, point to a possible homogenization of plant functional diversity and loss of ecosystems’ functional uniqueness, which would align with previously described global homogenization of species composition (Winter et al, 2009). Our results may also reflect a broader functional restructuring of ecological communities under human-driven environmental change. Previous work has shown that species occupying the “big, slow” end of trait space are disproportionately vulnerable to extinction in many taxa (Ripple et al, 2017; Carmona et al, 2021; Guo et al, 2026). Consistent with our findings for plants, Cooke et al (2019) projected that vertebrate assemblages will shift toward smaller, faster-lived, and more generalist species. If such patterns generalize across taxa, as our results suggest, ongoing human-mediated global change may increasingly reorganize ecosystems toward faster life-history strategies, favoring rapid turnover over long-term persistence.

The direction of the individual trait shift vectors appears to depend on a region’s initial position in trait space (Fig. 5). Previous studies have also found that there is a strong habitat dependency in what functional properties might make an introduced plant successful (Milanović et al, 2020; Wagner et al, 2017). In other words, functional dimensions that may be relevant for establishing a stable population in one type of habitat may not be in another. Our observations align with observations made regionally. In the United States Garbowski et al (2024) observed that in plant communities, along a gradient of introduced species abundance, community weighted means (CMWs) shifted towards smaller, faster, and more “do-it-yourself”-strategies, i.e., mycorrhiza-independence reflected in high specific root length. Studies focusing on Europe have shown that introduced species, especially those that become widespread and abundant, also tend to have higher SLA, but are taller, and have larger seeds (Küster et al, 2008; van Kleunen et al, 2010; Milanović et al, 2020). When translated to the level of whole ecosystem functional trait composition, this context dependency of change implies that the regional implications may also vary depending on which of the three dimensions of plant strategy is affected locally (Guo et al, 2022).

Although we use the term “global” to describe the scope of this study, we acknowledge the pronounced spatial bias in both CS and TRY data towards the Global North. In general, CS data are subject to well-known sampling biases, such as accessibility, seasonal sampling, or local population density (Meyer et al, 2016; Wolf et al, 2022; Mora et al, 2024; Tautenhahn et al, 2026). The shifts we detect here are thus possibly less sensitive to these general sampling biases, since they apply across all records. However, the species identification models that are integrated into most, if not all, citizen science species observation apps, are trained on images that are again biased toward European and North American plants. As mentioned above, European plants contribute disproportionately to the global introduced flora. This may introduce a bias in equatorial regions, where species observation tools may more reliably recognize introduced species than locally native floras.

Our results nonetheless indicate that regions underrepresented both in the data we use here and in the literature in general, may be most affected by changes in functional composition. Although the underlying samples are less complete and possibly more biased, we observe some of the strongest shifts along all three axes in Indomalaya and the Neotropics (Fig. 4). Previous work has not noted these regions to be particularity rich in introduced species (Pyšek et al, 2017), however our results suggest that on the level of plant functional composition, these regions seem to have been altered significantly through human-mediated plant species migration in the last centuries. In addition, many citizen scientists may not reliably distinguish between cultivated and naturally occurring plants (jharkness, 2020). While these factors could bias observations toward introduced species, they also mean that the CS occurrence data includes data on managed landscapes such as croplands, parks, and plantations. Consequently, our results might capture functional changes across both natural and human-modified ecosystems more reliably than studies restricted to so-called naturally occurring habitats, thus providing a broader perspective on shifts within trait space since the onset of the Anthropocene. In addition, the WCVP and GloNAF checklists may underestimate introduced species distributions due to their coarse resolution and potentially slow global updating, possibly making CS data a more realistic proxy in some regions. Nonetheless, it is important to note that these regions specifically contain comparatively few CS observations even though they harbor much of global plant diversity (Weigelt et al, 2020). The observed patterns may be confirmed or may change in the future as more data are collected. It is also important to note that our method has limited statistical power in small regions, such as islands, because too few grid cells are available to reliably detect significant differences.

A conceptual limitation of our study is that we assume the current relative abundances of native species observations broadly reflect historical baselines. In reality, introduced species may replace native species non-randomly, sometimes displacing functionally similar natives more than others (Liu et al, 2025b). If such selective replacement occurs, the trait shifts we observe could be somewhat exaggerated, but this caveat cannot be resolved with the data currently available. We therefore focus primarily on the direction of trait shifts and interpret the magnitude of shifts in trait space more cautiously.

Our approach captures shifts in functional space driven by introduced species at large scales. Although we cannot detect functional changes occurring exclusively within native-only assemblages, previous work has shown that introduced species play a disproportionate role in reshaping functional structure. Jaureguiberry et al (2022) showed that so-called invasive species are major drivers of shifts in species trait composition across taxa. In the absence of global historical baselines, leveraging introduced/native status therefore provides a pragmatic and scalable means of mapping functional change across a substantial fraction of human-altered terrestrial ecosystems.

In addition, introduced species tend to co-occur with broader anthropogenic pressures. While often framed as primary drivers of biodiversity loss, previous analyses indicate that they frequently emerge alongside other human pressures, such as grazing, land-use change, pollution, and the exploitation of natural resources (Jaureguiberry et al, 2022; Jauni et al, 2015). Consistently, the proportion of introduced species observations in the data visually aligns with cumulative anthropogenic pressures on biodiversity proposed by Bowler et al (2022), and exhibits a high correlation with human population density (*r* = 0.52, see Supplemental Fig. 7), mirroring previous findings of elevated numbers of introduced species in human-dominated landscapes (Richardson et al, 2025; Li et al, 2025).

Taken together, the strong correspondence between the proportion of introduced species observations in the citizen science data (Fig. 3) and previously mapped anthropogenic pressures on biodiversity (Bowler et al, 2022), combined with introduced species’ influence on trait composition (Jaureguiberry et al, 2022), suggests that our approach captures a broad, generalized signal of past shifts in global plant functional composition. While exceptions undoubtedly exist, it may be relatively uncommon for ecosystems to experience profound human-driven functional transformation without the establishment of introduced plant species. Thus, the trait shifts observed here might even be rather conservative, with the functional shifts contributed by introduced species representing only a baseline.

These broad-scale shifts in plant functional composition are likely to have had cascading consequences for ecosystem functioning. Because plant traits mediate the efficiency of carbon uptake, water use, and energy exchange with the environment (Reichstein et al, 2014; Gomarasca et al, 2023; Migliavacca et al, 2021), systematic shifts toward smaller, faster, shorter-lived plant assemblages may alter productivity dynamics and land–atmosphere interactions (Reichstein et al, 2014; Bonan, 2008). In this way, changes in functional trait composition extend beyond mere biodiversity patterns and likely influence ecosystem resilience and climate regulation (Mahecha et al, 2024; Westoby et al, 2002; Kambach et al, 2024; Lipoma et al, 2024; Sakschewski et al, 2016). Future research should explicitly link the large-scale shifts in functional trait space we present here to ecosystem functioning, coupled Earth system processes, including disturbance regimes, extreme event information, and multiple dimensions of resistance and resilience, in concert with other dimensions of biodiversity. The estimates of past trait distributions, may serve as input or constraints for ecosystem or Earth System models (Sakschewski et al, 2015; Rammer et al, 2024; Thonicke et al, 2020), which could shine light on the dynamics and implications of not only past, but also of future region-specific shifts in trait space.

## Methods

### Citizen science occurrences

We downloaded openly available citizen science (CS) occurrences of vascular plants (Tracheophyta) from GBIF on 15 June 2025 (GBIF, 2025) (*n* = 75,261,670). For the purpose of this paper, we define citizen science data specifically as large platform-based opportunistically sampled data by citizen scientists. Such CS data are subject to well-known sampling biases, such as accessibility, seasonal sampling, or urban areas (Meyer et al, 2016; Sierra et al, 2025; Tautenhahn et al, 2026). Datasets were manually selected from GBIF to include only opportunistically sampled, platform-based CS records (Quinn, 2025), so that the data would have more uniform sampling biases that are easier to interpret: iNaturalist Research-grade Observations (43 million records), Observation.org (17 million), Pl@ntNet (12 million), Vascular plant records verified via iRecord (1 million), Estonian Naturalists’ Society (1 million), Hatikka.fi (130,000), and Earth Guardians Weekly Feed (38,000). Occurrences close to botanical gardens, institutions, and country centroids, all ferns, lycopods and mosses were excluded using the CoordinateCleaner library in R (Zizka et al, 2019). The global sampling density is visualized in Supplemental Fig. 8.

### Trait data

Trait information was obtained from Joswig et al (2023), who predicted trait values using BHPMF gap-filling on TRY (Version 6) data (Kattge et al, 2020). The dataset contains estimates for 1, 056, 033 individual plants for 37 above- and below-ground traits (Table S1). Overall, 14% of values are missing, as the 10% of predictions with the highest uncertainty (STD) and phylogenetically inconsistent outliers were removed (Joswig et al, 2023). The compiled dataset was provided by the authors upon request; however, the original study provides reproducible code for generating the predictions from TRY data, which is openly available at www.try-db.org. We calculated species-level trait means for each species; about 50, 000 species in total.

### Native/introduced plant status

We retrieved the native or introduced status from the World Checklist of Vascular Plants Version 14 (WCVP) (Gov, 2025) and the Global Naturalized Alien Flora database Version 2.0 (GloNAF) (Davis et al, 2025). The resolution of distribution data in both datasets is TDWG level 3 (Brummitt, 2001), which corresponds mostly to individual countries and administrative borders at the sub-national level for larger countries. For each TDGW region, GloNAF lists all introduced species that have naturalized, i.e., became established a self-sustaining population. WCVP provides both native and introduced region polygons for each species.

To convert this data to the grid-cell level, we overlaid the polygons with an H3 hexagonal grid at resolution 3 (12,000 km^2^ on average per cell; Cooley and Shao, 2024) and added one buffer ring of hexagons to account for the coarse polygon boundaries, allowing us to assign species as native or non-native within each grid cell. We calculated a majority vote on the status (introduced/-native) for each grid cell based on the two datasets. If WCVP and GloNAF disagreed on native or introduced status, WCVP was prioritized. If there was a tie between statuses, due to overlapping region polygons, the species was labeled “native” for that grid cell. If there was no data available in either dataset on a species in the respective region, the occurrence was removed.

### Linking the different datasets

We merged the CS species observations with TRY species-level trait means via harmonized species names. We matched 95% of the CS vascular plant observations to species-level trait information. Taxonomic harmonization was based on the method detailed in Schellenberger Costa et al (2023). All scientific names were harmonized with WCVP as the main backbone. The status for each species and grid cell derived from WCVP and GloNAF was matched via harmonized species names and the respective h3 hexagon coordinate code. 1.4% of the occurrences were not matched to a status.

After filtering, taxonomic harmonization, and assigning native/introduced status, the dataset comprised *n* = 63,203,021 occurrences (*n*_native_ = 51,783,959; *n*_introduced_ = 11,419,062) across a total of 42, 520 species, of which 5, 521 had introduced status in at least one of the *xxx* hexagonal grid cells. All occurrences were matched to the 34 above- and 3 below-ground traits and assigned native/introduced labels. There was a total of 14% missing data in the occurrence *×* trait matrix.

### Past and present plant functional biogeography

Occurrence-level trait values were aggregated within each grid cell to calculate grid-cell means (H3, resolution 3 (Cooley and Shao, 2024)), a resolution that proved most robust when comparing CS-based trait maps to vegetation plot data (Wolf et al, 2022):

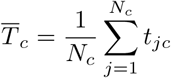

where *T̄_c_* is the mean trait value in grid cell *c*, *N_c_* is the total number of occurrences in grid cell *c*, and *t_jc_* is the trait value associated with occurrence *j* in grid cell *c*. Thus, species with more occurrences implicitly contribute more weight. Traits, such as leaf area, that have long-tailed distributions across species and differ orders of magnitude were log_10_-transformed before calculating the grid cell averages; see Table S1 for trait-specific details. This approach assumes that the grid cell means of traits derived from CS species observations are thus reflective of community weighted means (CWMs). Previous work showed that these two assumptions can be made, especially in well-sampled regions and when looking at broad, general patterns of grid-cell averages (Wolf et al, 2022; Lusk et al, 2026; Dechant et al, 2024).

Grid cell means *T̄_c_*were calculated for each trait and both time steps, i.e., past (native species occurrences only) and present (all available occurrence data for that grid cell). We filtered outliers based on the 0.01 and 0.99 quantile range.

### Dimension reduction

We calculated a principal component analysis (PCA) on the grid-cell means over all 37 traits. Both past and present grid-cell means for all traits were concatenated into one long matrix to calculate one overall PCA. The grid-cell averages *×* trait matrix had 6% missing data, and we used a mean imputation on the grid-cell means before calculating the PCA.

We chose not to filter grid cells by a minimum number of observations or trait coverage, as the number of records required to adequately represent functional trait space is likely to be region-specific. Consequently, the PCA includes substantial noise, which is reflected in the slow decay of variance explained across dimensions 4–37 (see Fig. 1d). Because it is unclear at what sampling intensity a grid cell reliably reflects local trait composition, we retained the unfiltered dataset and allowed sampling noise to be captured by higher-order dimensions to avoid introducing region-specific biases.

### Quantifying trait shifts

To summarize large-scale trait shifts, we grouped grid cells into biogeographic regions following biomes defined by Olson et al (2001) and further subdividing these biomes by continental realms. These biome definitions are based on expert knowledge and were obtained from the Terrestrial Ecoregions of the World dataset (www.worldwildlife.org/publications/terrestrial-ecoregions-of-the-world).

The statistical significance of regional trait shifts, including both individual traits and trait axes (PC1, PC2, and PC3), was assessed using unpaired Wilcoxon rank-sum tests comparing grid-cell mean values between past and present trait estimates. P-values were corrected for multiple testing using the less conservative Benjamini–Hochberg procedure to maintain sensitivity in this exploratory analysis, and shifts were considered significant at *α* = 0.05.

For each trait (including PC1, PC2, and PC3) and for each region, the magnitude of the difference between past and present trait distributions was quantified using Cliff’s delta (*δ*), equivalent to the rank-biserial correlation coefficient. Effect sizes were classified as negligible (*|δ| <* 0.11), small (0.11 *≤ |δ| <* 0.28), medium (0.28 *≤ |δ| <* 0.43), or large (*|δ| ≥* 0.43) following Peres (2025). Regions with p-values *<* 0.05 and non-negligible Cliff’s *δ*, i.e. *|δ| ≥* 0.11, were selected as regions with most prominent shifts and are marked in Figures 5a.

Grid-cell differences in trait space (Fig. 4) were calculated for each principal component (PC1, PC2, and PC3) as Δ*PC_c_* = *PC_c,_*_present_ *− PC_c,_*_past_, where *PC_c_* is the PC value for grid cell *c*. Regional centroid shift vectors were defined in 3D PCA space, with the vector origin at the mean of past grid-cell PCA scores and the vector endpoint at the mean of present grid-cell PCA scores for each region. Projections of these vectors into 2D PC1/PC2 and PC3/PC2 space are shown in Fig. 5b–c.

## Data Availability

All data used to produce the results presented here are openly available (though some require creating a user account and preprocessing): Citizen science data download (GBIF, 2025), TRY Version 6.0 and gap-filling work-flow (Kattge et al, 2020; Joswig et al, 2023), Global Naturalized Alien Flora database Version 2.0 (GloNAF) (Davis et al, 2025), World Checklist of Vascular Plants Version 14 (WCVP) (Gov, 2025), human population density at 1° (Center for International Earth Science Information Network (CIESIN) - Columbia University, 2017).

## Code Availability

We provide a fully reproducible workflow here: (https://github.com/sojwolf/ Anthropocene Shifts Trait Space).

## Use of Artificial Intelligence

LLMs (GPT-4 and GPT-5) functioned as an interactive research assistant – as a discussion partner and a technical aide. LMMs did not access raw data, run analyses, or generate final results. All code was written (using GitHub-Copilot–chat and tab-complete–for AI-assisted coding), executed, tested, and validated by the authors. Scientific interpretations and methodological choices were made by the authors. A small handful of papers we cite here were found by the AI academic literature research tool Elicit. A detailed report of the use of LLMs can be found in the Supplemental Materials 14.

## Acknowledgments

This study was funded by the National Research Data Infrastructure Germany for Biodiversity, NFDI4Biodiversity, a project by the German Research Foundation (DFG), project number 442032008, and by the European Space Agency Climate Change Initiative (ESA-CCI) Tipping Elements SIRENE project (contract no. 4000146954/24/I-LR). The study is supported by the TRY initiative on plant traits (http://www.try-db.org). T.K. acknowledges funding by DFG within the project PANOPS (project no. 504978936) and by ESA FORTRACK via the ESA CLIMATE SPACE: Climate-Biodiversity studies. D.S. acknowledges funding from the Philipp Schwartz-Initiative of the Alexander von Humboldt-Stiftung (grant no. 232201751). K.M. acknowledges funding by the Saxon State Ministry for Science, Culture and Tourism (SMWK)—(3-7304/35/6-2021/48880) and by ESA AI4Science project ‘Deep-Features’ 2024–2026. L.C. acknowledge the German Research Foundation (DFG) funding via iDiv (DFG FZT 118, 202548816). P.P. was supported by long-term research development project RVO 67985939 (Czech Academy of Sciences). S.W. thanks David Reher for discussions on statistical methodology and Negin Katal and Daniel Lusk for discussions on traits and figures, and Jan Pergl for insight on introduced species data. We thank the vegetation scientists, who measured plant traits, digitized them, or made them available in databases. Last, but certainly not least, we thank all the citizen scientists who volunteer their time and expertise to build the many datasets collected in GBIF.

## Author Contributions

S.W., D.S. and T.K. conceived the study, S.W. performed the analyses and wrote the original draft. D.S. and D.S.C. performed data preprocessing. T.K., M.D.M., and C.W. supervised the project. S.W., D.S., M.W., G.K., K.M., K.N., S.T. and L.C. developed the methodology. M.K., P.W., H.K., and P.P. curated data. All authors contributed to writing and editing the manuscript.

## Supplementary information

**Table 1:** List of plant functional traits and their units.

| TraitID | TraitName | UnitName |
| --- | --- | --- |
| 4 | Stem specific density (SSD) or wood density | g/cm<sup>3</sup> |
| 6 | Root rooting depth | m |
| 13 | Leaf carbon (C) content per leaf dry mass | mg/g |
| 14 | Leaf nitrogen (N) content per leaf dry mass | mg/g |
| 15 | Leaf phosphorus (P) content per leaf dry mass | mg/g |
| 21 | Stem diameter | m |
| 26 | Seed dry mass | mg |
| 27 | Seed length | mm |
| 46 | Leaf thickness | mm |
| 47 | Leaf dry matter content (LDMC) | g/g |
| 50 | Leaf nitrogen (N) content per leaf area | g m<sup>-2</sup> |
| 55 | Leaf dry mass (single leaf) | mg |
| 78 | Leaf nitrogen isotope signature ($\delta^{15}\text{N}$) | ‰ |
| 95 | Seed germination rate | % |
| 138 | Seed number per reproduction unit | number |
| 144 | Leaf length | mm |
| 145 | Leaf width | cm |
| 146 | Leaf C/N ratio | g g<sup>-1</sup> |
| 163 | Leaf fresh mass | g |
| 169 | Stem conduit density | mm<sup>-2</sup> |
| 223 | Chromosome number | n |
| 224 | Chromosome cDNA content | pg |
| 237 | Dispersal unit length | mm |
| 281 | Stem conduit diameter | μm |
| 282 | Wood vessel element length | μm |
| 289 | Wood fiber lengths | μm |
| 297 | Wood ray density | mm<sup>-1</sup> |
| 351 | Seed number per dispersal unit | – |
| 614 | Specific fine root length (SRL) | cm/g |
| 1080 | Specific root length (SRL) | cm/g |
| 3106 | Plant height (vegetative) | m |
| 3107 | Plant height (generative) | m |
| 3112 | Leaf area (petiole undefined) | mm<sup>2</sup> |
| 3113 | Leaflet area (petiole undefined) | mm<sup>2</sup> |
| 3114 | Leaf/leaflet area (undefined) | mm<sup>2</sup> |
| 3117 | SLA (petiole undefined) | mm<sup>2</sup> mg<sup>-1</sup> |
| 3120 | Leaf water content (dry mass basis) | g/g |

**Fig. 6:**
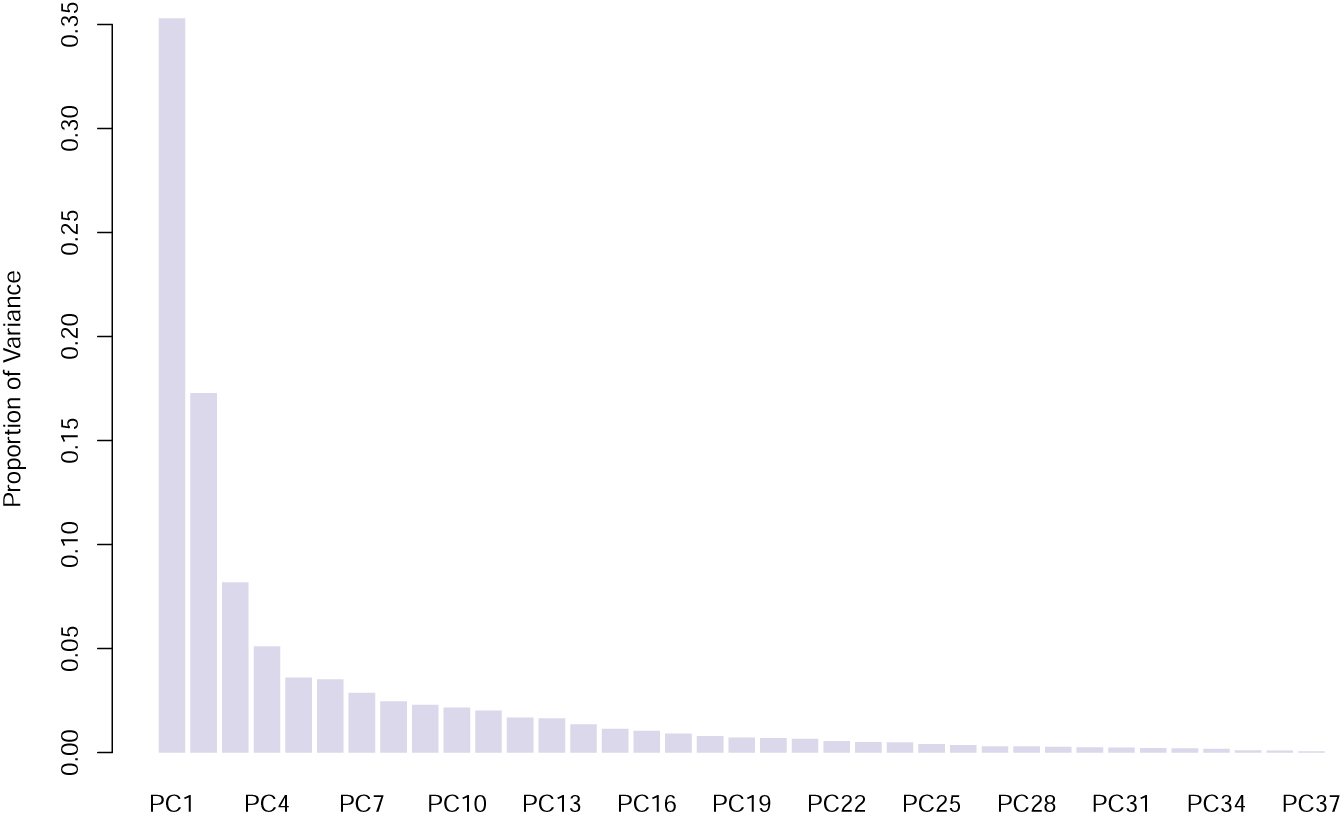
Proportion of variance explained by each principal component

**Fig. 7:**
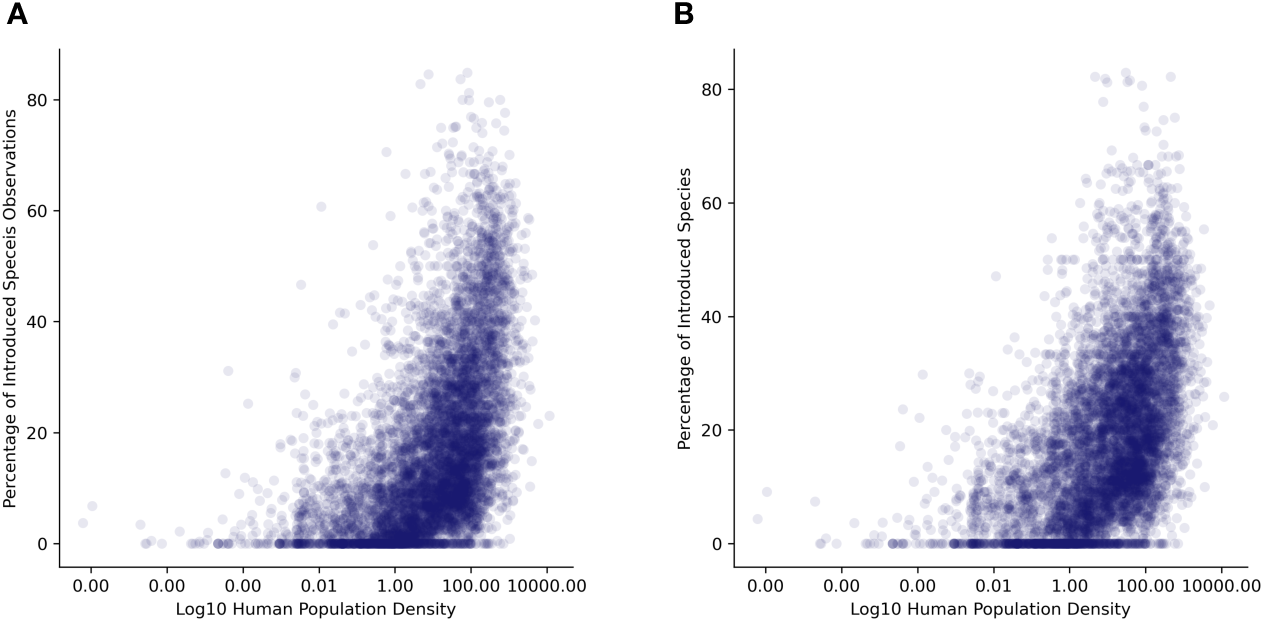
Scatter plots showing relationship between human population density (log10) and **A** percentage of introduced species observations, *r* = 0.52; and **B** percentage of introduced species, *r* = 0.50.

**Fig. 8:**
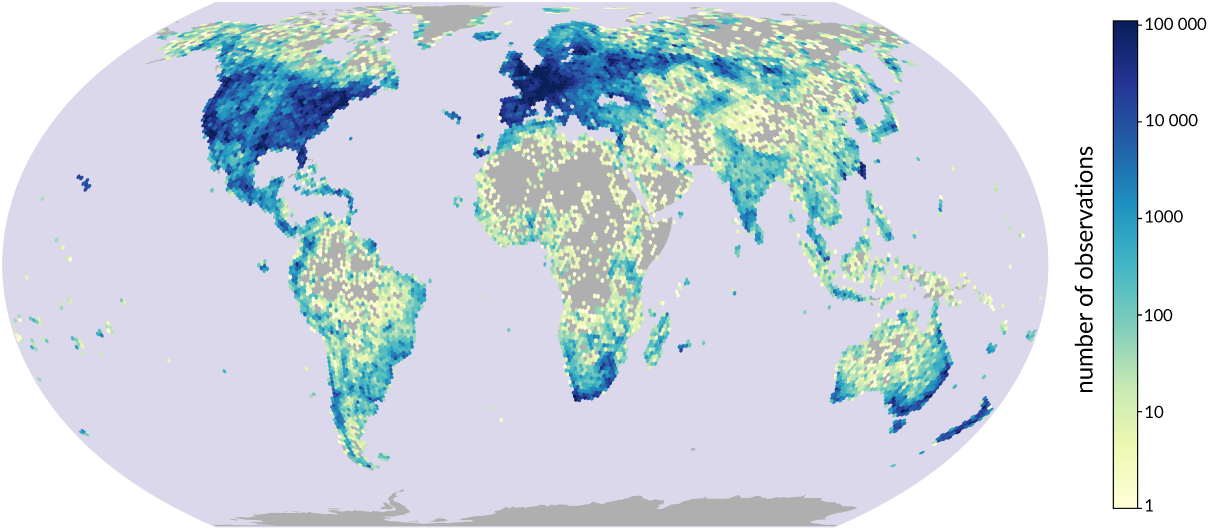
Distribution and density of citizen science occurrence data, both native and introduced species occurrences (number of observations *log*10-transformed)

**Fig. 9:**
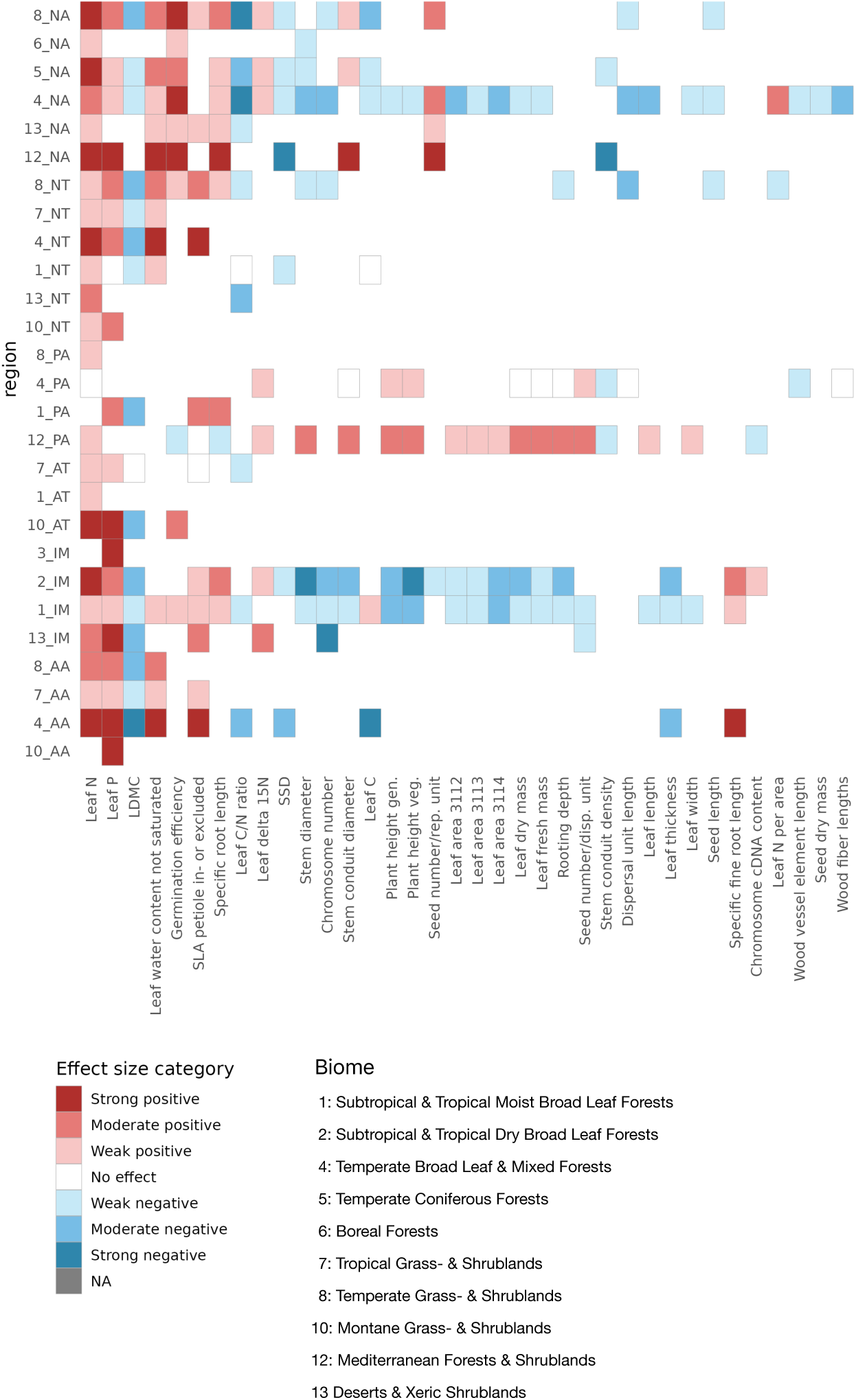
Effect sizes of individual trait shifts per region with adjusted p-value *<* 0.05

**Fig. 10:**
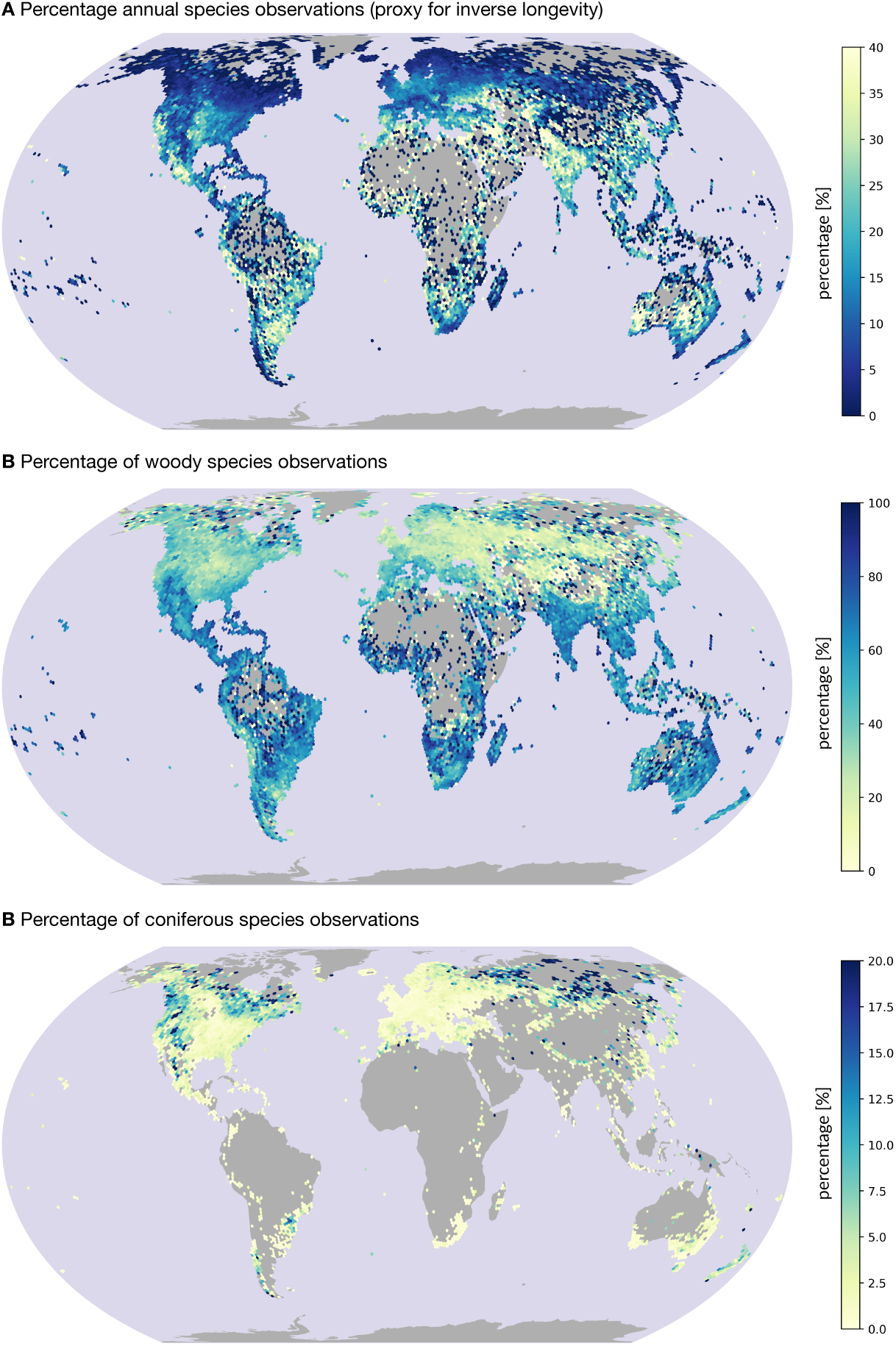
**A** Percentage of annual species occurrences per grid cell **B** Percentage of woody species occurrences per grid cell**C** Percentage of conifer species occurrences per grid cell

### Use of Large Language Models

**Note**: This section was generated using the following prompt in ChatGPT 5.0: “Could you list all our past discussions about the trait shift science project and your contribution? I want to make a transparent part in the supplement showing how I used LLMs. Feel free to be detailed and don’t understate your role.”

Output:

#### General Role of the LLM

ChatGPT (OpenAI) was used as an interactive research assistant throughout the trait-shift project. Its role extended beyond copy-editing or syntactic code assistance and included conceptual discussion of analytical strategies, methodological design, iterative refinement of data structures and workflows, debugging and reasoning about geospatial and trait-based joins, and support for scaling, optimization, and reproducibility.

The LLM did not access raw data, execute analyses, or make autonomous scientific decisions. All data processing, analyses, and scientific interpretations were implemented, validated, and finalized by the author.

#### Conceptual framing of trait shifts

Discussions focused on how to operationalize “trait shifts” in a global invasion context. The author defined the core scientific question concerning trait differences between native and global (“all”) occurrence groups and selected the relevant trait and environmental dimensions. The LLM contributed by helping articulate alternative conceptual framings of trait shifts, discussing within-cell versus across-cell comparisons, and serving as a sounding board for aligning ecological interpretation with data structure.

#### Integration of occurrence data with trait datasets

The author curated and cleaned GBIF-based occurrence data and linked these records to species-level trait information. The LLM assisted in reasoning about join strategies, potential pitfalls of trait sparsity and missing data, and the biological implications of aggregating traits across occurrences.

#### Trait data structure and variable handling

The project involved managing datasets containing multiple trait variables, derived variables, grouping factors, and spatial indices. The LLM contributed by suggesting programmatic approaches for selecting and manipulating trait columns, advising on reproducible data structures, and supporting the decision to treat PCA axes as traits in downstream analyses.

#### Use and interpretation of PCA

Principal component analysis was used to summarize multivariate trait variation. The author computed the PCA and determined the biological interpretation of the resulting axes. The LLM supported discussion of the conceptual implications of using PCA axes as response variables, comparability across groups, and limitations that should be explicitly acknowledged in the manuscript.

#### Spatial aggregation using discrete global grids

Trait shifts were spatially analyzed using discrete global grid systems (H3). The author selected the spatial resolution and implemented the spatial subsetting. The LLM contributed by explaining the conceptual properties of H3 grids, comparing them to alternative spatial frameworks, and discussing resolution trade-offs and spatial autocorrelation.

#### Group-wise trait comparisons within spatial units

The author designed and implemented comparisons of trait values between “native” and “all” groups within spatial grid cells. The LLM assisted in conceptualizing within-cell comparisons, discussing statistical implications of unequal sample sizes, and exploring appropriate ways to summarize and visualize trait shifts.

#### Computational scaling and feasibility

Given the global scale of the dataset, computational feasibility was a recurring topic. The author optimized pipelines and made decisions about data reduction and aggregation. The LLM contributed suggestions for scaling strategies, identification of steps where dimensionality reduction was appropriate, and balancing ecological resolution with computational constraints.

#### Parallel processing design

Parallelization strategies were discussed to accelerate spatial and trait-based workflows. The author implemented multiprocessing and managed merging of outputs. The LLM contributed to reasoning about how trait calculations interact with parallelization, trade-offs between parallelizing over space versus observations, and potential inefficiencies or failure points.

#### Visualization strategy

The author produced all figures and selected final visualization approaches. The LLM contributed ideas for grid-based summaries and trait contrasts, assisted in evaluating interpretability for readers, and supported alignment between figures and the core research questions.

#### Reproducibility and transparency

Ensuring reproducibility and methodological transparency was an explicit goal of the project. The author structured scripts, data flow, and documentation. The LLM assisted in articulating pipeline logic, identifying steps that warranted explicit documentation, and contributing to the drafting of transparent descriptions of LLM use.

#### Summary of LLM Contribution

In this project, the LLM functioned as a methodological co-thinker, a workflow and code-logic assistant, a conceptual sounding board for trait-based and spatial ecological analyses, and a writing support tool for transparency and documentation. The LLM meaningfully shaped how analyses were reasoned about and structured, but did not generate results, run analyses, or interpret outcomes independently of the author.

